# Network based statistics reveals trophic and neuroprotective effect of early high dose erythropoetin on brain connectivity in very preterm infants

**DOI:** 10.1101/533901

**Authors:** András Jakab, Christoph Rüegger, Hans Ulrich Bucher, Malek Makki, Petra Hüppi, Ruth Tuura, Cornelia Hagmann, the Swiss EPO Neuroprotection Trial Group

## Abstract

Periventricular white matter injury is common in very preterm infants and it is associated with long term neurodevelopmental impairments. While evidence supports the protective effects of erythropoetin (EPO) in preventing injury, we currently lack the complete understanding of how EPO affects the emergence and maturation of anatomical brain connectivity and function. In this case-control study, connectomic analysis based on diffusion MRI tractography was applied to evaluate the effect of early high-dose EPO in preterm infants. A whole brain, network-level analysis revealed a sub-network of anatomical brain connections in which connectivity strengths were significantly stronger in the EPO group. This distributed network comprised connections predominantly in the frontal and temporal lobe bilaterally, and the effect of EPO was focused on the peripheral and feeder connections. EPO resulted in a globally increased clustering coefficient and higher local efficiency, while higher strength, efficiency and increased clustering was found for regions in the temporal lobe, supramarginal gyrus, inferior frontal gyrus, in the caudate and cingulate gyri. The connectivity network most affected by the EPO treatment showed a steeper increase in FA with age compared to the placebo group. These results demonstrate a weak but widespread effect of EPO on the structural connectivity network and a possible trophic effect of EPO reflected by increasing network segregation, predominantly in local connections.

## Introduction

While the mortality rate of very preterm infants has decreased significantly within the last decade (Ruegger et al., 2014), long-term neurodevelopmental problems such as cognitive, motor and behavior impairments remain of concern (Latal, 2009; Saigal and Doyle, 2008). Periventricular white matter injury (PWMI) is the predominant form of perinatal brain injury in preterm infants PWMI is characterized by marked astrogliosis and microgliosis, and initially by a decrease in premyelinating oligodendrocytes (pre-OLs) (Haynes et al., 2003). PWMI is accompanied by neuronal/axonal deficits in the cerebral white matter, thalamus, basal ganglia, cerebral cortex, brainstem and cerebellum. This led to the term "encephalopathy of prematurity“, which describes a complex amalgam of primary destructive disease and secondary maturational and trophic disturbances (Volpe, 2009).

Encephalopathy of prematurity has been linked to later neurodevelopmental impairments such as cerebral palsy, motor dysfunction, cognitive and behavioral problems as well as deficits in executive functions (Woodward et al., 2006; Woodward et al., 2011). The high incidence of neurodevelopmental impairments in preterm infants is reflected by the ongoing search for neuroprotective interventions that can prevent brain injury or enhance repair of the immature brain, with the ultimate goal of improving long-term motor and cognitive outcome (Gonzalez et al., 2009).

Among several pharmacological candidates to prevent brain injury or improve its development in preterm infants, erythropoietin (EPO) has been shown to be among the most promising neuroprotective agents (Robertson et al., 2012). Protective effects of EPO important for reducing acute injury include decreased apoptosis, inflammation, excitotoxicity and glutamate toxicity (Juul, 2012; Shingo et al., 2001). Furthermore, EPO was shown to play a role in developmental mechanisms, as it stimulates neurogenesis (i.e. proliferation and differentiation of pre-oligodendrocytes (Sugawa et al., 2002)), angiogenesis and migration of regenerating neurons (Tsai et al., 2006). In 2012, recruitment of the first randomized, double-blind placebo-controlled, prospective multicentre trial (NCT00413946) in very preterm infants using high-dose EPO as a potential neuroprotective agent was completed (n=450 preterm infants). In a subgroup of infants, MR imaging showed less PWMI on conventional MRI (Leuchter et al., 2014), and improved white matter microstructure assessed by tract based spatial statistics (O’Gorman et al., 2015).

Given the widespread brain changes observed in these prior studies and the clear relationship of global brain networks and brain function (van den Heuvel and Sporns, 2013), it is of interest to examine the global brain network connectivity in these infants. In recent years, due to improvements of MR imaging such as in diffusion MRI, it has become possible to map the global brain connectivity enabling us to study the global network organizational properties of the brain, the effect of neuroprotective interventions and their relation to brain development and pathologies (Hagmann et al., 2012).

The complete structural or functional organization of the brain is represented by its connectome – the brain-network level architecture of all neuronal connections (Sporns, 2011). Numerous studies using dMRI proved that it is possible to map the developing connectome of infants with a granularity that is comparable to adult studies (Brown et al., 2014; Cao et al., 2017; Jakab et al., 2015; Tymofiyeva et al., 2012; van den Heuvel et al., 2015). We are only beginning to understand if neurodevelopment in congenital or acquired diseases can lead to a global impairment of the interactions between the neural elements, and whether this is reflected by impairments in the macro-scale connectome. A recent study using connectome analysis has demonstrated specific alterations in brain topology and structural organization in the brain network organization of high-risk preterm born children at school age and such altered regional brain connectivity was related to specific neurocognitive deficits in children born preterm (Fischi-Gomez et al., 2015; Fischi-Gomez et al., 2016).

Hence, connectomic analysis might not only help to further understand the neural correlate of neurodevelopmental impairments in preterm infants but also to help understand the effects of EPO on global brain connectivity development.

Our study is built on two hypotheses. First, we hypothesize that the neuroprotective effect manifests as changes in the brain’s structural connections, which may only become apparent on the whole-brain network level and not locally. Connectomic analysis may therefore shed light on the effects of therapeutic agents not only on isolated brain regions, but also on how they affect the topography of brain connections (Bonilha et al., 2015; Crossley et al., 2017; Zeng et al., 2015). Second but most importantly, EPO treatment may steer the ongoing structural connectivity development into an alternative, more optimal trajectory, which would in turn be observed as global changes in the structural connectivity architecture compared to untreated subjects.

## Methods

Ethical approval was granted by the local ethical committee (KEK StV36/04), and the study was approved by the Swiss drug surveillance unit (Swissmedic, 2005DR3179). The trial was registered at ClinicalTrials.gov (number NCT00413946).

### Patient population

The inclusion criteria and selection process for the study have been described in detail previously (Leuchter et al., 2014; Natalucci et al., 2016; O’Gorman et al., 2015). In short, the preterm infants in this study represent a subgroup of infants enrolled in the randomized, double-blind placebo-controlled, prospective multicentre study “Does erythropoietin improve outcome in preterm infants” (NCT00413946). For this analysis, only infants with DTI performed in Zurich (n=140) were included. Clinical co-morbidities such as sepsis (blood culture proven), NEC, PDA, and CLD (defined as oxygen requirement at corrected 36 weeks of gestation) were noted. Our database comprised the socio-economic status (SES) and corrected gestational age at the time of MRI (CGA). Neurodevelopment at two years of age was characterized by the mental developmental index (MDI) according to the Bayley Scales of Infant and Toddler Development, Second Edition (Bayley-II).

### Randomization, neuroprotective intervention and blinding

Two year neurodevelopmental outcomes and the treatment protocol have been has been previously published (Fauchere et al., 2008; Natalucci et al., 2016). Study medication was randomly assigned to each patient number in a 1:1 allocation using a computer-based random-number generator. Erythropoetin or an equivalent volume of normal saline (NaCl 0.9%) placebo was administered intravenously before 3 hours of age after birth, at 12-18 and at 36-42 hours after birth. A single dose consists of 25 μg (3000 IU) human erythropoietin per kilogram body weight dissolved in 1 ml sterile water. Hospital staff, parents, the outcome assessors and data analysts were kept blinded to the allocation.

### Cerebral MRI protocol

Cerebral MRI was performed with a 3.0 T GE scanner (GE Medical Systems), using an 8-channel receive-only head coil. All infants were scanned under natural sleep using a vacuum mattress. Ear plugs (attenuation: 24 dB; Earsoft; Aearo) and Minimuffs (attenuation: 7 dB; Natus) were applied for noise protection. Oxygen saturation was monitored during scanning, and a neonatologist and a neonatal nurse were present during the MRI investigation.

We acquired T1-weighted MR images with a 3D fast spoiled gradient echo sequence (echo time = 2.6 ms, repetition time = 5.7 ms, inversion time = 750 ms, flip angle = 12°, voxel resolution 0.7 × 0.7 × 1.4 mm3), and T2-weighted images with a fast recovery fast spin echo sequence (echo time = 126 ms, repetition time = 6600 ms, resolution 0.7 × 0.7 × 1.5 mm3).

DTI was acquired using a pulsed gradient spin echo echo-planar imaging sequence with echo time = 77 ms, repetition time = 9 s, field of view = 18 cm, matrix = 128 × 128, slice thickness = 3 mm. The diffusion encoding scheme included 21 non-collinear gradient encoding directions with b = 1000 and four interleaved b = 0 images. Of the 140 infants scanned in Zurich, in 78 infants DTI data demonstrated motion artefacts (n=78), 4 infants had with cystic lesions (n=4) apparent on structural MRI and hence were excluded from further analysis, resulting in a final group size of 58 infants, of whom 24 were treated with EPO and 34 were treated with placebo.

### Diffusion tensor image processing

DTI data were visually controlled for artifacts by observing each diffusion gradient volume and each image slice for signal dephasing or spin history artifacts caused by movements. If the newborn woke up or moved excessively during the DTI scan, data were discarded and the subject was excluded. Subjects were also excluded if more than 3 diffusion-weighting gradient volumes were corrupted by motion artifacts. Image frames and the corresponding entries in the b-matrix and b-value descriptor files were removed from further analysis if head movement caused extensive signal dropout throughout the brain in the given frame. A custom script (**Digital Supplement**) written in bash programming language for Linux was used to process the neonatal DTI images. Spurious image shifts originating from eddy currents were corrected with the *eddy_correct* command and the b-matrix was reoriented with the *fdt_rotate_bvecs* script in the FSL software. Next, image intensity shifts along the supero-inferior direction were corrected by normalizing the intensity of each axial slice to the corresponding slice in the motion-corrected, median diffusion-weighted image. Rician noise filtering of DTI was performed using a command line tool in Slicer 3D (Aja-Fernandez et al., 2009). FA and MD maps were calculated by running the *dtifit* command in FSL on the Rician noise filtered data.

### Region of interest system

Anatomical parcellation of the brains was carried out by using the neonatal regions of interest (ROI) from UNC Infant atlas UNC (Shi et al., 2011). This comprised 90 cortical and subcortical areas that were propagated from the Automated Anatomical Labeling (AAL: (Tzourio-Mazoyer et al., 2002)) to the images of 33.4 – 42.1 gestational week neonates (for ROI nomenclature, see Table S1). We first created a custom fractional anisotropy template in the UNC space, which was used to transfer the UNC-AAL ROIs to the DTI space of each subject.

This FA template was constructed in three steps based on DTI data of a separate study population, consisting of 40 normally developing and term-born newborns imaged within 6 weeks after birth (CGA of the subjects: 42.5 ± 1.9 (39 – 48.7) weeks, 20 females and 20 males). First, we registered the B_0_ images of each neonate with the T2-weighted template of the UNC atlas using an affine registration followed by an elastic non-linear registration implemented in the NIFTIreg software (reg_f3d command, control grid size: 9 * 9 * 9 mm, weight of the bending energy penalty term: 0.05, gradient smoothing with a kernel of 4 mm). The transformations arising from this registration step were used to re-sample the Rician filtered FA images of each neonate to UNC template. An initial FA template was then created by averaging these FA images. Each subject’s FA maps were co-registered to the initial FA template with NIFTIreg with an elastic deformation allowing for finer distortions (reg_f3d command, control grid size: 5 * 5 * 5 mm, weight of the bending energy penalty term: 0.005, gradient smoothing with a kernel of 4 mm).

### Structural connectivity network construction

Seed points for probabilistic diffusion tractography (Probabilistic index of connectivity method (Parker et al., 2003) were defined as the voxels with FA ≥ 0.1 within a whole-brain mask. Orientation density functions (ODFs) were estimated using fourth order Spherical Harmonics and a maximum of two local ODF maxima were set to be detected at each voxel and probability density function (PDF) profile was produced from the local ODF maxima. Fiber tracking was carried out on the voxel-wise PDF profile with the Euler interpolation method using 10 iterations per each seed point. Tracing stopped at any voxel whose FA was less than 0.2. The tractography script is available as digital supplement.

Two kinds of weighted, undirected structural connectivity networks were formed based on whole-brain probabilistic tractography: FA_mean_ and SC. In both networks, nodes corresponded to the AAL ROIs in subject space. In the FA_mean_ structural connectivity network, the strength of each edge was given by the mean fractional anisotropy (FA) value of all tractography streamlines connecting the regions at the two endpoints.

Secondly, network edges were defined using the normalized structural connectivity strength, SC. Tractography streamline counts connecting any two nodes were normalized for the bias arising from the volumetric differences between the regions of interests corresponding to the nodes, and for the linear bias that arises from more distant brain regions showing more streamlines (Hagmann et al., 2008).

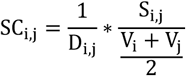

Where SC_i,j_ is the structural connectivity strength, D_i,j_ is the Euclidean distance between the node centre-points, S_i,j_ is the number of tractography streamlines connecting nodes i and j, V_i_ and V_j_ are the node volumes.

The control for the presence of spurious connections, the FA_mean_ network was restricted to edges with FA_mean_ value larger than 0.2. False positives were controlled for the SC network by keeping the 40% strongest connections on the individual and on the group level (de Reus and van den Heuvel, 2013).

## Graph theoretical analysis

Structural brain network analysis comprised connection (edge) selection with an iterative thresholding procedure. First, structural connectivity (SC) networks were thresholded from a density of 1% to 99%, and the same surviving edges were kept for both the FA_mean_ and SC networks. For both networks at each cost-thresholded level, the following graph theoretical parameters were calculated using the Brain Connectivity Toolbox for Matlab (Rubinov and Sporns, 2010).

**Strength** of region *i*:

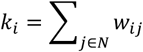

*where w*_*ij*_ *is the connection weight between i and j, N is the entire network.*

**Betweenness centrality** of region *i* (Freeman, 1979):

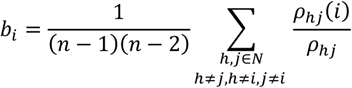

*where* ρ_*hj*_ *is the number of shortest paths between h and j, and* ρ_*hj*_*(i) is the number of shortest paths between h and j that pass through i. N are all network nodes. h and j are the neighboring nodes of i.*

**Global efficiency** of the weighted network N:

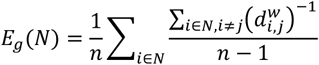

*where n is the total number of nodes in the network N, 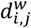 is the shortest weighted path length between nodes i and j.*

Weighted **local efficiency** is the efficiency calculated for a sub-network constructed from the 1^st^ neighbors of the node i:

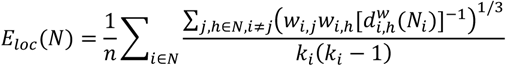

Where *w*_*i,j*_, *w*_*i,h*_ are connection weights between nodes i,j and i,h, 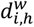 *is the shortest weighted path length between nodes i and h, k*_*i*_ *is the degree of node i.*

**Average local efficiency** is the average of all local efficiencies across the network nodes.

**Weighted clustering coefficient** of the network (Onnela et al., 2005; Watts and Strogatz, 1998):

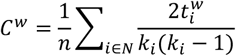

*where 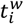 is the weighted geometric mean of triangles around node i:*

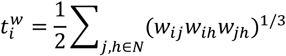

## Core and periphery structure

The connectivity networks were subdivided into two non-overlapping groups of nodes, a core and a periphery based on the Kernighan-Lin algorithm for graph partitioning (Borgatti and Everett, 2000; Newman, 2006), implemented as the “core_periphery_dir” command in the Brain Connectivity Toolbox for Matlab. A common core for the study population was defined as the nodes that were in the core partition in more than 90% of the subjects. Core connections connected the core nodes, their first degree connections were defined as feeders, while the remaining non-zero connections were labeled as peripheral connections. For each subject, the core-ness parameter was calculated that reflected the degree of separation between the core and peripheral nodes (Borgatti and Everett, 2000; Newman, 2006).

## Statistical analysis

To test the effects of EPO treatment and neurodevelopment on the global and nodal graph theoretical parameters, a multivariate analysis of covariance in Matlab R2014 using the mancovan toolbox (Mathworks Inc., Mattick, USA) was performed. During the mass-multivariate testing of graph theoretical parameters across network nodes (brain regions) statistical significance was adjusted for multiple comparisons using the Benjamini-Hochberg procedure, implemented in the FDR toolbox in Matlab R2014. As networks were tested at various cost thresholds, only those results were accepted that survived FDR adjustment across nodes at more than two cost threshold levels between 20% and 90%, and consistently showed the same effect direction (positive or negative group difference) in this cost range.

Hypothesis tests on structural connectivity networks were based on the network-based statistics (NBS) method described by Zalesky et al. (Zalesky et al., 2010), which was implemented in the NBS Toolbox for Matlab R2014. NBS avoids the problem of multiple comparisons during mass univariate tests on connectivity networks by estimating statistical significance for subsets of mutually connected network nodes in topological rather than physical space (Reess et al., 2016). The NBS method comprises four steps. First, the t-statistic for each individual edge in the connectivity network is calculated. Second, a primary component-forming threshold (p<0.05, uncorrected) is applied to identify edges displaying differences in connectivity strength. Third, subthreshold edges are assessed for mutual connections forming clusters in topological space that may point toward the existence of non-chance clusters. Fourth, a test with 5000 random permutations is applied to compute statistical significance for all previously identified network component. As NBS uses permutation test to build up the sample distribution, it can be applied to smaller study groups without assuming normality. The final hypothesis test is then carried out for the empirically determined components by comparing their extent with the proportion of permutations yielding a component with equal or greater size, correcting for the family-wise error rate at cluster level with p<0.05. In our tests, a primary threshold t-score of 2 was chosen. Next, a post hoc analysis was performed using linear regression in SPSS V22 (IBM, Armonk, New York) to select any further demographic or clinical parameters that may have an effect on the structural connectivity values averaged over the networks from the NBS analysis.

We presented the results of the NBS as three-dimensional graph visualizations, which represented below p-threshold connection pairs surviving multiple comparison correction. The brain networks were visualized with the BrainNet Viewer for Matlab R2014 (Xia et al., 2013).

## Results

### Patients

58 preterm infants with mean (SD) corrected gestational age at birth 29.75 (1.44) weeks and at scanning of 41.1 (2.09) weeks) were included into the connectomic analysis. **Table 1** shows the demographic and clinical details of the two groups. No significant differences were found in CGA at scanning or in any other neonatal or clinical details between the two treatment groups. Subjects with cystic lesions were not included in the current study. Based on the white matter injury scoring system of Woodward et al. (Woodward et al., 2006), 54 (93.2%) neonates displayed normal white matter and only 4 (6.8%) had abnormal white matter.

**Table 1.** Demographic and clinical parameters of the infants. Differences of clinical characteristics between groups were evaluated with the chi-square test or the Fisher exact test as appropriate for the categorical variables, and with Student’s t test or Mann-Whitney test as appropriate for the continuous variables. SD – standard deviation, IQR – inter-quartile range (25^th^-75^th^ percentiles). ^+^ sepsis proven by positive blood cultures; NEC, necrotizing enterocolitis, ** oxygen requirement at corrected age of 36 weeks.

| Characteristic | Reported results | EPO group (N=24) | Placebo group (N=34) | P-value |
| --- | --- | --- | --- | --- |
| Gestational age (weeks) | mean ± SD | 30.17 ± 1.44 | 29.5 ± 1.44 | .095<sup>2</sup> |
| Birth weight (g) | mean ± SD | 1337 ± 332 | 1192 ± 10 | .098<sup>2</sup> |
| Z-score of birth weight* (SD) | mean ± SD | -0.27 ± 0.71 | -0.19 ± 0.95 | 0.72<sup>2</sup> |
| Gestational age at MRI (weeks) | mean ± SD | 40.93 (2.09) | 41.33 (2.12) | 0.49<sup>2</sup> |
| Gender (female) | N (%) | 8 (33%) | 14 (41%) | 0.61<sup>1</sup> |
| Retinopathy of prematurity | N (%) | 3 (13%) | 4 (12%) | 0.77<sup>1</sup> |
| Sepsis* | N (%) | 3 (13%) | 5 (15%) | 0.88<sup>1</sup> |
| NEC | N (%) | 0 (0%) | 1 (3%) | 0.58<sup>1</sup> |
| Mechanical ventilation | N (%) | 8 (33%) | 13 (38%) | 0.46<sup>1</sup> |
| Duration of mechanical ventilation (days) | median (range) | 0 (0; 6) | 0 (2; 12) | 0.73<sup>3</sup> |
| Bronchopulmonary dysplasia** | N (%) | 2 (8%) | 5 (14%) | 0.38<sup>1</sup> |
| Postnatal steroids | N (%) | 2 (8%) | 0 (0%) | 0.17<sup>1</sup> |

<sup>1</sup> Chi-square test or Fisher exact test as appropriate
<sup>2</sup> Student test
<sup>3</sup> Mann-Whitney test

### Quality of neonatal DTI data

In the EPO group, corrupted gradient volumes were excluded from the analysis in two subjects (1 and 2 volumes removed). 1-3 volumes (1.57 ± 0.8) were removed for 7 subjects in the placebo group. Network level analysis revealed that the number of DTI frames removed did not correlate with the FA_mean_ or SC values (highest significance, NBS, p=0.422 and p=0.131, respectively).

### Global graph theory characteristics

Global efficiency, average local efficiency, average clustering coefficient and average nodal strength were depicted as a function of network density separately for the EPO and placebo groups in **Fig. 2/a,c,e,g**. EPO treated newborns showed 5.2% higher average clustering coefficient, with maximum effect observed at a network density of 27% (**Fig. 2/a**, T=7.79, p=0.0072, estimated marginal means (EMM) EPO: 0.077 (CI, 95%: 0.075 – 0.079), Placebo: 0.073 (CI, 95%: 0.072 – 0.075), appearing at CGA=41.2 weeks).

**Figure 1.**
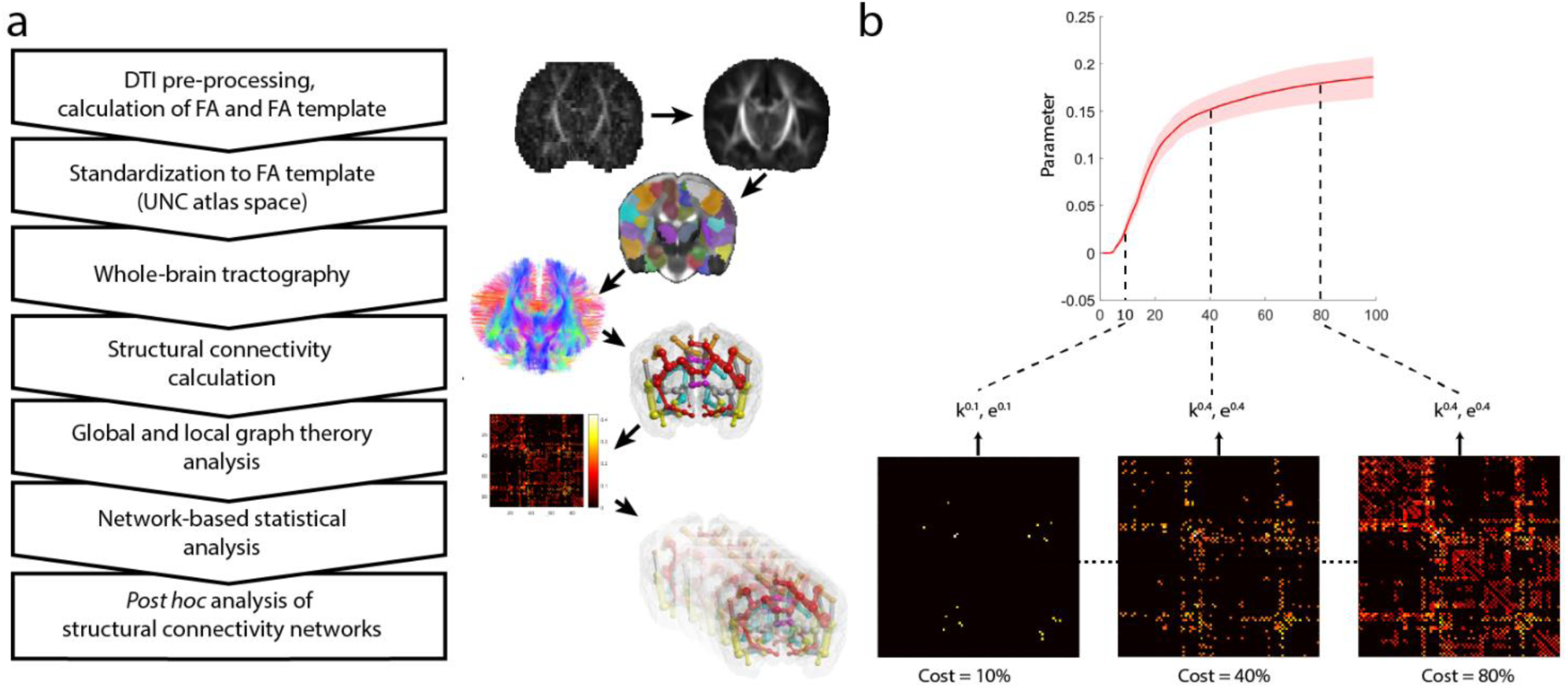
Methodological overview. (a) image post processing and connectivity analysis steps, (b) thresholding procedure based on network density (cost).

**Figure 2.**
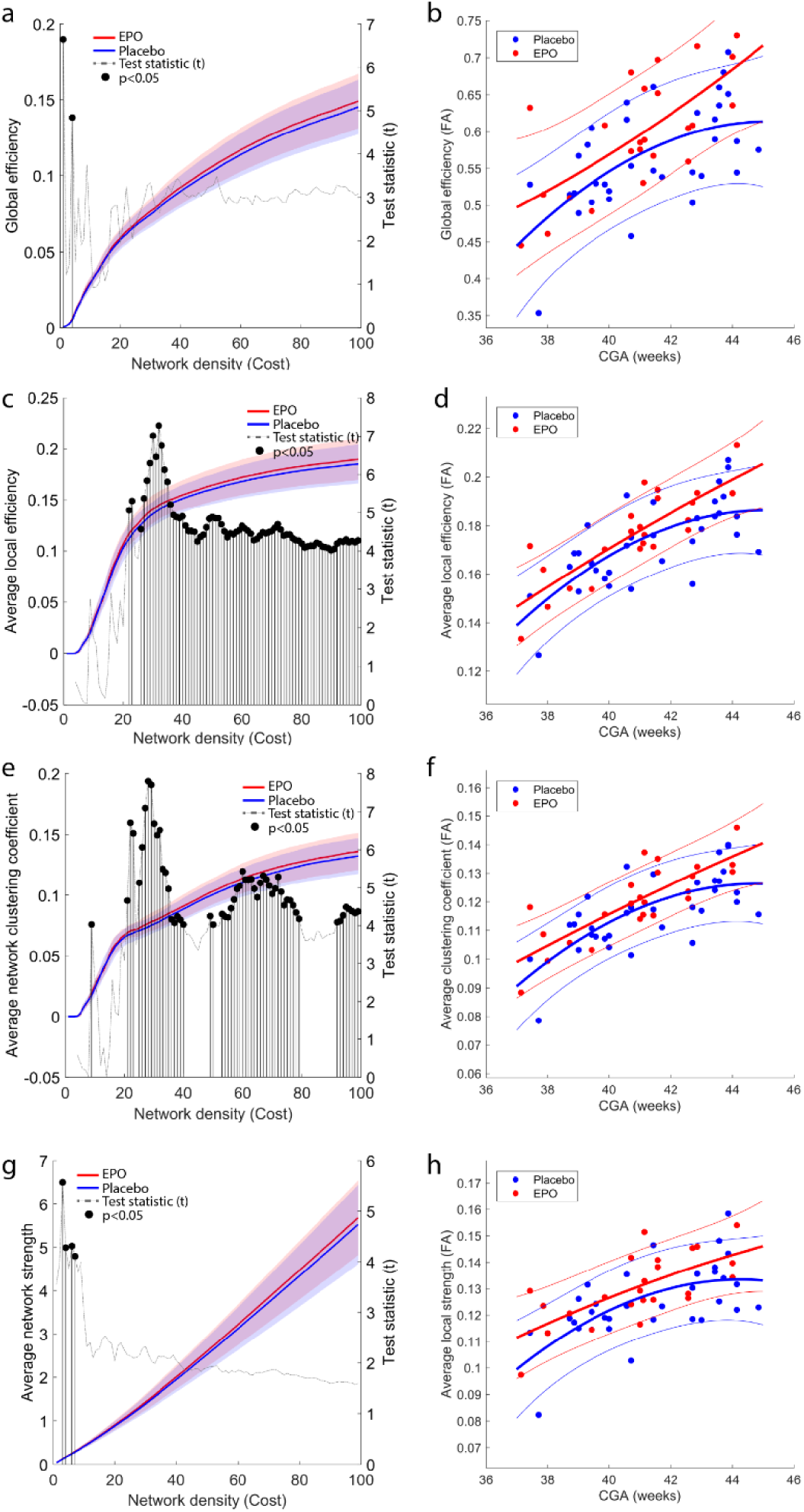
Global graph theory characteristics of the EPO and placebo treated groups. (a,c,e,g) global characteristics are plotted (left axis) as a function of network density. The right axis shows the T statistic of the multivariate analysis of group differences. Network costs at which significant EPO-placebo differences appear are marked with segmented line and dot (p<0.05, FDR-corrected across network nodes). Blue lines: mean values, placebo group, red lines: mean values, EPO group, transparent area reflects the variability of measurements (± 1SD range). Network density threshold is expressed as a percentile of total nonzero connections. (b,d,f,h) the network characteristics that were significantly different across group are plotted as a function of CGA.

Average clustering coefficient increased with CGA (**Fig. 2/c**), and the steepness of this was moderately larger after the 42. week in the EPO group (**Fig. 2/d**). Global network efficiency and average nodal strength was not affected by EPO treatment in the density range that we used to define statistical significance (**Fig. 2/a,b and Fig. 2/g,h**). Average local efficiency was 4.3% higher in the EPO group, maximum effect was observed at a network density of 31% (**Fig. 2/e**, T=7.28, p=0.0093, EMM, EPO: 0.146 (CI, 95%: 0.142 – 0.150), Placebo: 0.140 (CI, 95%: 0.137 – 0.143), appearing at CGA=41.2 weeks), the age dependency showed similar tendency to the clustering coefficient (**Fig. 2/f**).

Mental developmental index at 2 years of age was not correlated with the global graph theoretical findings that were affected by EPO treatment (average clustering coefficient, p=0.095, average local efficiency, p=187, multivariate model adjusted for parental SES and CGA).

### Local graph theory characteristics (Figure 3)

Local graph theory characteristics of six nodes in the FA_mean_ network were significantly different between the EPO and placebo groups, the results of the nodal analysis are summarized in **Table 2**. Graph theoretical measures of three temporal brain regions were significantly different between the EPO-treated and placebo groups. The treatment appeared to have a predilection towards nodes that were part of the network periphery, the only core network node affected by treatment was the right caudate of the SC connectivity network.

**Table 2.**
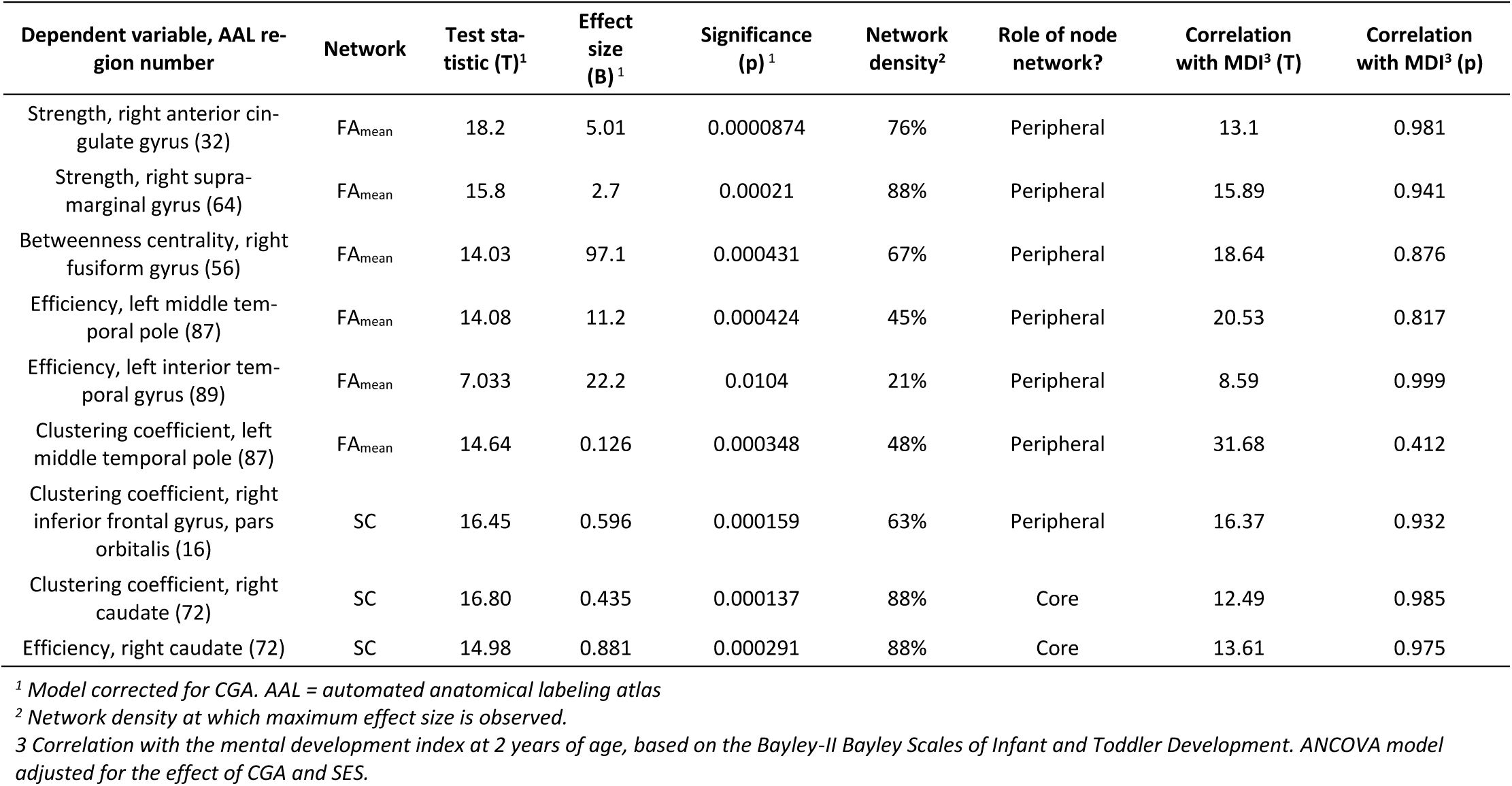
Effect of EPO treatment – local graph theory characteristics.

**Figure 3.**
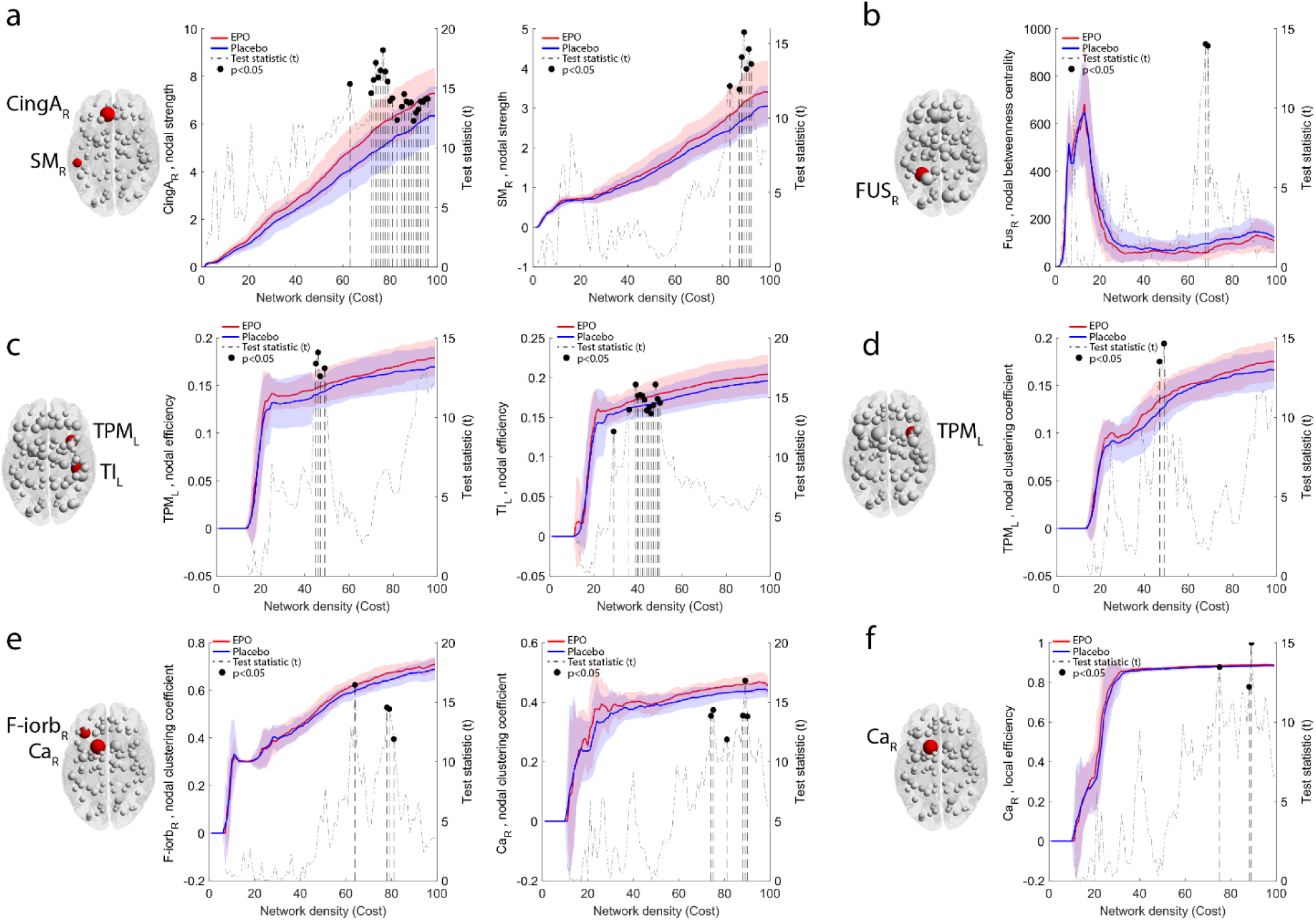
Effect of EPO treatment on nodal graph theory characteristics. (a) nodal strength of the right anterior cingulate and supramarginal gyri, (b) nodal betweenness centrality of the right fusiform gyrus, (c) nodal efficiency of the left medial temporal pole and inferior temporal gyrus, (d) nodal clustering coefficient of the left middle temporal pole, (e) SC-network, nodal clustering coefficient of the right frontal infraorbital gyrus and the caudate, (f) local efficiency of the right caudate. Nodal characteristics are plotted (left axis) as a function of network density, while the right axis shows the T statistic of the multivariate analysis of group differences. Network density threshold is expressed as a percentile of total non-zero connections. Network costs at which significant EPO-placebo differences appear are marked with segmented line and dot (p<0.05, FDR-corrected across network nodes). Blue lines: mean values, placebo group, red lines: mean values, EPO group, transparent area reflects the variability of measurements (± 1SD range).

In the FA_mean_ network, the left middle temporal pole showed 10.3% higher clustering coefficient in the EPO-treated neonates (p=0.00348, EMM, EPO: 0.139 (CI, 95%: 0.134 – 0.145), placebo: 0.126 (CI, 95%: 0.122 – 0.131). The local efficiency of the left middle temporal pole and left inferior temporal gyrus was 7.14% and 12.6% higher in EPO treated newborns, respectively, calculated based on the estimated marginal means of the multivariate model (left middle temporal pole, p=0.000424, EMM, EPO: 0.15 (CI, 95%: 0.146 – 0.154), placebo: 0.14 (CI, 95%: 0.137 – 0.143), left inferior temporal gyrus, p=0.0104, EMM, EPO: 0.161 (CI, 95%: 0.148 – 0.175), placebo: 0.143 (CI, 95%: 0.131 – 0.154)). At a network density of 67%, the betweenness centrality of the right fusiform gyrus was 41.3% lower in the EPO treated group (p=0.000431, EMM, EPO: 57.07 (CI, 95%: 40.719 – 73.42), placebo: 97.07 (CI, 95%: 83.34 – 110.79). The strengths of the right anterior cingulate gyrus and right supramarginal gyri were 21.8% and 18.4% higher in EPO (right anterior cingulate gyrus, p=0.0000874, EMM, EPO: 6.01 (CI, 95%: 5.707 – 6.491), Placebo: 5.07 (CI, 95%: 4.678 – 5.34), right supramarginal gyrus, p=0.00021, EMM, EPO: 3.187 (CI, 95%: 2.996 – 3.378), placebo: 2.692 (CI, 95%: 2.532 – 2.852)). Three graph theoretical parameters of two nodes were found to be affected by EPO treatment in the SC network. The clustering coefficient of the right inferior frontal gyrus pars orbitalis and right caudate was 5.2% and 6.2% higher in EPO treated newborns, respectively, (right inferior frontal gyrus pars orbitalis, p=0.000159, EMM, EPO: 0.627 (CI, 95%: 0.615 – 0.638), placebo: 0.596 (CI, 95%: 0.586 – 0.605), right caudate, p=0.000137, EMM, EPO: 0.462 (CI, 95%: 0.452 – 0.472), placebo: 0.435 (CI, 95%: 0.426 – 0.443)). In the SC network, the right caudate, which is part of the core network, showed 0.68% higher efficiency in the EPO-treated neonates (p=0.000291, EMM, EPO: 0.887 (CI, 95%: 0.885 – 0.890), placebo: 0.881 (CI, 95%: 0.879 – 0.883). Mental developmental index at 2 years of age was not correlated with any of the nodal graph theoretical findings that were affected by EPO treatment (**Table 2**).

### Effect of gestational age on the structural connectivity networks

Structural connectivity strength increased with the corrected gestational age (CGA) of the neonates. FA_mean_ values were found to be significantly positively correlated with CGA in 455 of the **1362** nonzero edges (33.4%) of the structural connectivity network (**Fig. S1**). SC increased with CGA in 322 network edges (23.6%) of the structural connectivity network (**Supplementary Fig. S1**). These results prompted us to correct the following statistical analysis steps for the effect of CGA.

### Core structure and network based statistics

We found that 39 (2.86% of 1362 non-zero network edges) connections in the FA_mean_ network (chosen network density: 35%) were significantly stronger in the neonates treated with EPO when controlling for the effect of CGA. The biggest difference was measured for intra-lobar medium range, frontal, limbic, temporal and occipital connections (**Fig. 4/a**). The structural connections within this network were characterized by 10.2 % higher FA_mean_ in the EPO group than in the placebo-treated neonates (p= 0.000089, estimated marginal means (EMM) from a linear regression model controlling for the effect of CGA, EPO group: 0.195 (CI, 95%: 0.188 – 0.202), Placebo group: 0.177 (CI 95%: 0.171 – 0.182), appearing at CGA=41.2 weeks). The network nodes with the highest number of significantly different network edges (>3 significant edges) were the right anterior and middle cingulate gyri, left and right superior frontal gyri, right middle temporal gyrus and left middle occipital gyrus (**Fig. 4/a**). The mean network strength of the connections affected by EPO treatment were not correlated with the mental developmental index at 2 years of age (T=8.587, p=0.999).

**Figure 4.**
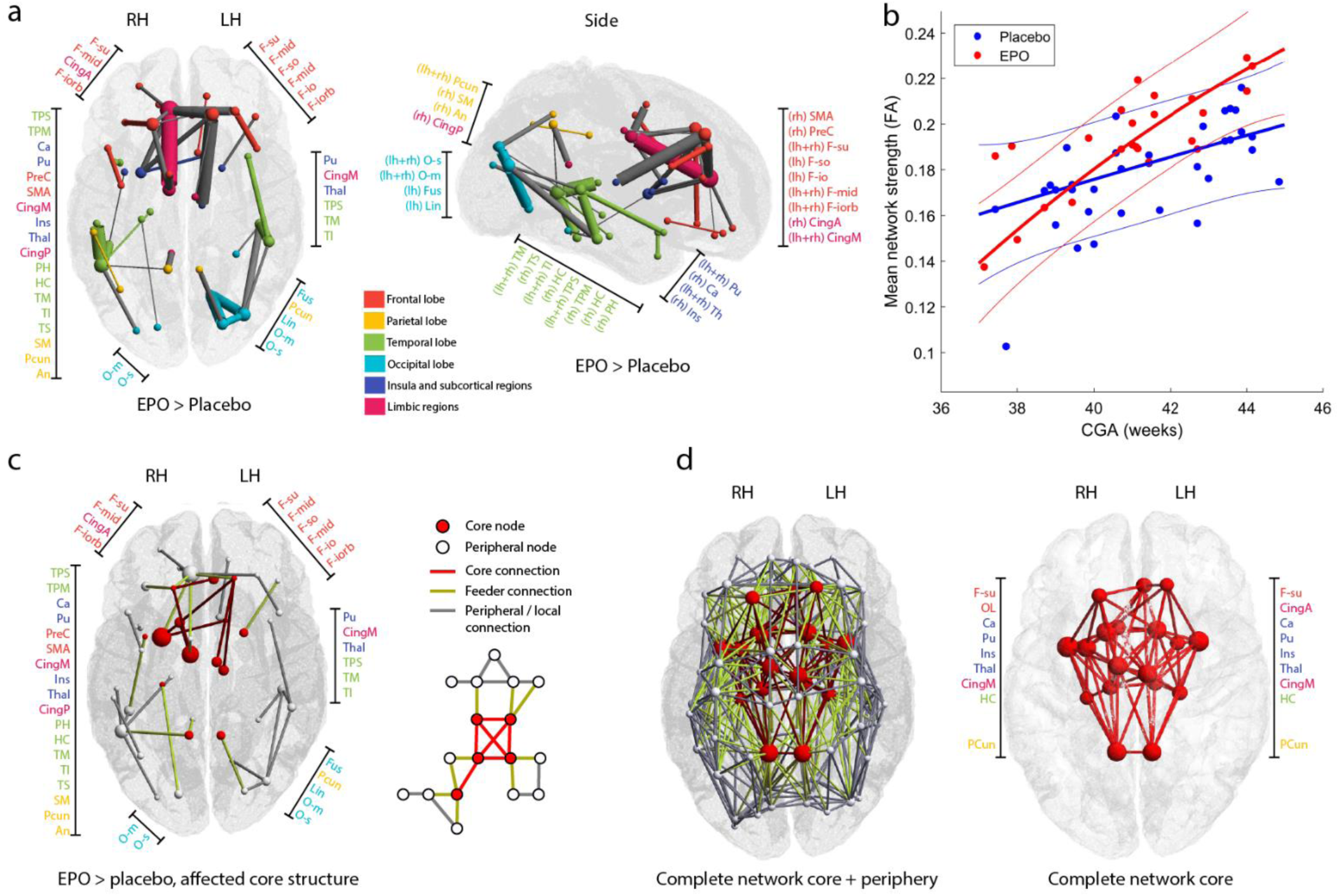
Effect of EPO treatment: network based statistics and core structure analysis. (a) structural connectivity network (FA_mean_) with increased connection strength in EPO-treated neonates. The size of nodes correlates with the number of significantly positive edges arising from the node, while edge width correlates with the test statistic of the NBS analysis. Gray edges mean inter-modular, colored edges intra-modular or intra-lobar connections. LH: left hemisphere, RH: right hemisphere. (b) linear regression plot showing the relationship between average FA_mean_ of the significant graph edges, CGA at the time of MRI and group allocation. (c) network core affected by EPO treatment, NBS results, (d) common network core of the study population. In the middle image, core and peripheral nodes are depicted in red and yellow, respectively, while core connections, feeder connections and local connections are depicted as red, yellow and gray lines. For the abbreviations of ROIs, see Table S1.

Next, we reconstructed the core connectivity structure in the study population. The core consisted of the 9-9 hemispherically symmetric nodes (**Fig. 4/c,d**): superior frontal gyrus, anterior and middle cingulate gyri, caudate, putamen, thalamus, insula, hippocampus and precuneus, which largely overlaps with the previously reported rich-club structural core (van den Heuvel and Sporns, 2011). Out of the 39 network edges affected by EPO treatment, 7 were core connections (17.9% of network, connections: anterior cingulate – superior frontal gyrus, middle cingulate – superior frontal gyrus bilaterally, putamen – superior frontal gyrus, thalamus – superior frontal gyrus middle cingulate - caudate, and caudate - putamen), 12 were feeders (30.8%) while the rest were peripheral (51.3%) connections. The coreness statistic was not significantly different between EPO and placebo-treated new-borns (T=2.213, p=0.1425). The affected core network structure is illustrated in **Fig. 4/c**.

At a density of 35%, NBS revealed no differences between the EPO and placebo groups when using the SC connectivity network instead of the FA_mean_.

As the edge strength correlated with CGA across the whole structural connectivity network (**Supplementary Fig. S1**) and the steepness of the regression line appeared to differ between placebo and EPO treated groups **(Fig. 4/b**), we tested whether group difference varied as a function of CGA (Group-CGA interaction).

A two-group NBS analysis with continuous covariate interaction was performed, which revealed that interaction is not significant in any of the network edges in the FA_mean_ connectivity network (minimum p=0.089). Next, we repeated the statistical test to capture the effects of EPO treatment on the structural connectivity networks, this model was corrected for the Group-CGA interaction term. The effects of EPO on the FA_mean_ connectivity network were seen in an almost identical set of edges to the previously performed tests with covariate interaction. With covariate interaction, 92.5% of the edges were overlapping (Dice coefficient of significant edges) with the tests not accounting for covariate interaction in the FA_mean_ network. The effects of EPO with and without Group-CGA interaction are demonstrated in **Supplementary Fig. S2**.

## Discussion

In this study, we applied connectomic analysis to identify the effect of early high dose EPO administration on the structural brain connectivity network in preterm infants. While we have previously shown that early high dose EPO administration reduces brain injury and improves white matter development in the major white matter tracts (O’Gorman et al., 2015), little is known about the effects of EPO on connectivity networks in the preterm brain.

Our findings reflect an early, weak effect of the EPO treatment on the overall structural brain connectivity of preterm infants. This effect was not limited to any of the large anatomical subdivisions, such as lobes of the brain, but the connectome of EPO treated infants was rather characterized by a widespread increase of local structural connectivity strengths. Global network strength – the sum of connectivity weights of all connections of a node – was not affected by the EPO treatment. The increased average local efficiency, statistically unchanged global efficiency after EPO treatment reflects the better ability of the connectivity network to transmit information at the local level and a shift towards a more regular network. We examined four local topological properties of the structural brain networks: network integration was measured by nodal strength and efficiency, while segregation was indicated by the clustering coefficient and betweenness centrality (Onnela et al., 2005).

It has been shown that brain hubs form a so-called “rich club” or core structure (van den Heuvel and Sporns, 2011), which is characterized by a tendency for high-degree nodes to be more densely interconnected. This structure is likely present as early as the 30^th^ gestational week (Ball et al., 2014), and during neurodevelopment after very preterm birth, this core is most likely prioritized over local connectivity (Fischi-Gomez et al., 2016; Karolis et al., 2016). In a recent study, a stronger rich-club architecture was found in preterms than in normally developing infants, despite a relative deficiency of white matter resources (Karolis et al., 2016). Our network based statistical analysis revealed that EPO facilitates both the development of peripheral and frontal, thalamic and limbic core connections. The EPO-induced development of peripheral, i.e., local connectivity, may provide valuable means to counteract the structural connectivity impairments that are specific for preterm birth (Karolis et al., 2016).

Our results reflect a weak, but widespread, local effect in the structural connectivity network after EPO treatment, which may give rise to more concentrated changes in the role of frontal, temporal and subcortical regions in the entire network. These regions are presumed to be densely interconnected areas – hubs – early on. Clustering and the formation of hubs is an emergent feature of the human connectome. Clustering coefficient is a measure of brain network segregation and is used to quantitatively describe the brain’s ability for specialized processing within such interconnected groups of brain regions (Sporns, 2013).

We found increased clustering coefficient in the left inferior temporal gyrus, middle temporal pole and inferior frontal gyrus. The clustering coefficient of the caudate, which is part of the preterm core connectivity network, was increased after EPO treatment. An increase in this metric is the direct consequence of a higher prevalence of clustered connectivity around a brain region (Watts and Strogatz, 1998), which may potentially reflect increased anatomical and functional segregation of this area (Rubinov and Sporns, 2010). Results from neuroimaging studies imply that the majority of large-scale pathways are already present by birth (Doria et al., 2010; van den Heuvel et al., 2015), and the newborn connectome features an adult-like smallworld modular architecture that could be sensitive to disruptions, as shown, for example, in children after hypoxic ischemic encephalopathy (Tymofiyeva et al., 2012). van den Heuvel el al. found that clustering coefficient averaged over the entire structural connectivity network increased from the 30th gestational week until term equivalent age in preterm infants (van den Heuvel et al., 2015), which is in line with our observations of gradually increasing clustering coefficients. Increased anatomical segregation may support more efficient consolidation of information from different functional networks through long-range connections, and it is safe to assume that during neurotypical development, clustering coefficient of the most important connectivity hubs gradually increase (van den Heuvel and Sporns, 2013). The developmental predilection for white matter injury to occur during prematurity appears to relate to both timing of appearance and regional distribution of susceptible pre-oligoendrocytes (Back et al., 2007; Riddle et al., 2006). The deep white matter myelinates in a caudal-to-rostral, proximal to distal, central to peripheral direction, with sensory pathways myelinating before motor pathways and with deep occipital white matter maturing first and frontal white matter maturing last (Kinney et al., 1988). Hence, the increased clustering coefficients in the frontal and temporal white matter in EPO treated infants might also reflect a regional and temporal variability of the EPO effect. Postnatal age has been associated with increasing connectivity in the frontal areas (Pandit et al., 2014). Our results suggest that EPO might enhance the normal development of connections with increasing age in predominantly frontal areas. This is an important finding as previous imaging studies have shown that FA in the frontal white matter lobe was reduced by prematurity (Rose et al., 2008). Such white matter alterations seem to persist into adolescence and adulthood (Vollmer et al., 2017; Yung et al., 2007), and such alterations were correlated with measures of executive function and general cognitive abilities in adolescence and young adulthood (Vollmer et al., 2017).

Based on previous works (Dosenbach et al., 2007), we propose that the newborn and infant brain gradually builds up the ability of its connectivity network to integrate information between its subparts. During this process, the presumed neuroprotective effect of EPO could steer the development into a trajectory that is characterized by a steeper increase of clustering coefficients. The most prominent feature of the developing infant brain in dMRI is the rapid increase of the diffusion anisotropy. We found that the corrected gestational age at MRI correlates with mean FA in many network edges after correcting for multiple comparisons, which is most likely the consequence of the progression of myelination (Berman et al., 2005; Huppi et al., 1998). Fractional anisotropy is known to increase gradually during the last trimester of gestation, and this trajectory is not significantly different in the postnatal development of prematurely born infants (Bonifacio et al., 2010). We found that the connectivity network most affected by the EPO treatment showed a steeper increase compared to the placebo group, however, no significant group-CGA interaction was found. A similar tendency was observed for the clustering coefficient, efficiency and strength during the nodal graph theory analysis. The increasing FA is considered to reflect an increase in white matter structure and thus an increase in the efficacy of long-distance neuronal signal transmission in the neonatal brain (van den Heuvel et al., 2015). The described steeper increase in FA and in clustering coefficient might reflect the long-term effects of EPO on neurogenesis (Juul, 2010). Based on these findings, we may also speculate that a possible trophic effect of EPO manifests as increasing network integration, especially at more advanced CGA.

A recent meta-analysis including 6163 children born very preterm and 5471 term-born children showed that children born very preterm have lower IQs, lower scores on measures of executive functioning and lower scores on measures of processing speed (Brydges et al., 2018). Recent neuroimaging and lesion studies suggest that executive functions depend on distributed networks encompassing both frontal and posterior (mainly parietal) associative cortices (Collette et al., 2006; Jurado and Rosselli, 2007). This is consistent with an imaging study of children born very preterm, in which frontal network alterations have been associated with higher-cognitive function deficits (Fischi-Gomez et al., 2015; Fischi-Gomez et al., 2016). A recent meta-analysis showed that prophylactic EPO improved cognitive outcome at 18 months of infants born very preterm (Fischer et al., 2017). Whether the improved structural connectivity in the EPO treated children seen in this study is reflected by better higher cognitive function such as executive function has to be shown.

We identified the following technical limitations in our study protocol. The low number of diffusion-weighting directions (n=21) allows estimating only one tensor per image voxel reliably; this inherently affects the result of probabilistic tracking of connections. Newborns underwent MRI in natural sleep, and longer scan time during DTI will greatly increase the ratio of corrupted image frames due to head motion, and this limits the number of applicable diffusion-weighting directions. During pre-processing, we used linear registration to correct eddy-current induced distortions, during which residual mis-alignment can adversely impact model fitting to the data. Improved correction would be achieved by techniques that model the diffusion signal and correct for eddy-currents and head motion in one resampling step (Andersson and Sotiropoulos, 2016). More accurate modeling of the diffusion propagator function and the application of more complex tractography techniques to the newborn brain would result in more reproducible description of the structural connectome in the future (Kuhn et al., 2015; Lebel et al., 2012; Wakana et al., 2007). We found only a weak effect of the EPO treatment on the overall structural brain connectivity of preterm infants. This might be explained by the timing and short duration of the EPO intervention.

Early high dose administration of EPO in very preterm infants shows a neuroprotective and trophic effect on the global brain connectivity, which effect was found to be marginally larger at a more advanced infant age. The connectome of EPO treated infants was characterized by increased average local efficiency, and increased structural connectivity strengths in edges that contribute to the peripheral connectivity in the presence of a preserved connectivity core. This increased integrity and local efficiency of the structural brain connectome, however, was not reflected as improved mental or psychomotor development at two years of age, therefore the beneficial effects of EPO treatment on neurodevelopment might manifest during later life.

## Funding

This work was supported by the OPO Foundation, the FZK Foundation, the “Stiftung für wissenschaftliche Forschung an der UZH” and the EMDO Foundation, R.T. was supported by the EMDO Foundation. The study was supported by a grant received from the Swiss National Science Foundation (SNF 3200B0-108176).

### The Swiss EPO Neuroprotection Trial Group

The following local investigators and hospitals participated in this study (study sites are listed in alphabetical order): Aarau: Kinderklinik Kantonsspital Aarau (Georg Zeilinger, MD; Sylviane Pasquier, MD); Basel: Universitätskinderklinik UKBB (Christoph Bührer, MD; René Glanzmann, MD; Sven Schulzke, MD); Chur: Abteilung für Neonatologie, Kantons-und Regionalspital (Brigitte Scharrer, MD; Walter Bär, MD); Geneva: Division of Development and Growth, Department of Pediatrics (S. Sizonenko MD); Neonatology Unit, Department of Pediatrics (Riccardo Pfister, MD); Geneva CIBM: F. Lazeyras Zürich: UniversitätsSpital Zürich, Department of Neonatology (Jean-Claude Fauchère, MD, Brigitte Koller); Zürich: Centre of MR Research (Martin Ernst, MD; Hadwig Speckbacher, MTRA); Zurich Pharmacy: (D. Fetz, B. Christen).

### This manuscript is the author’s original version

## Supplementary Materials

**Supplementary Figure S1.**
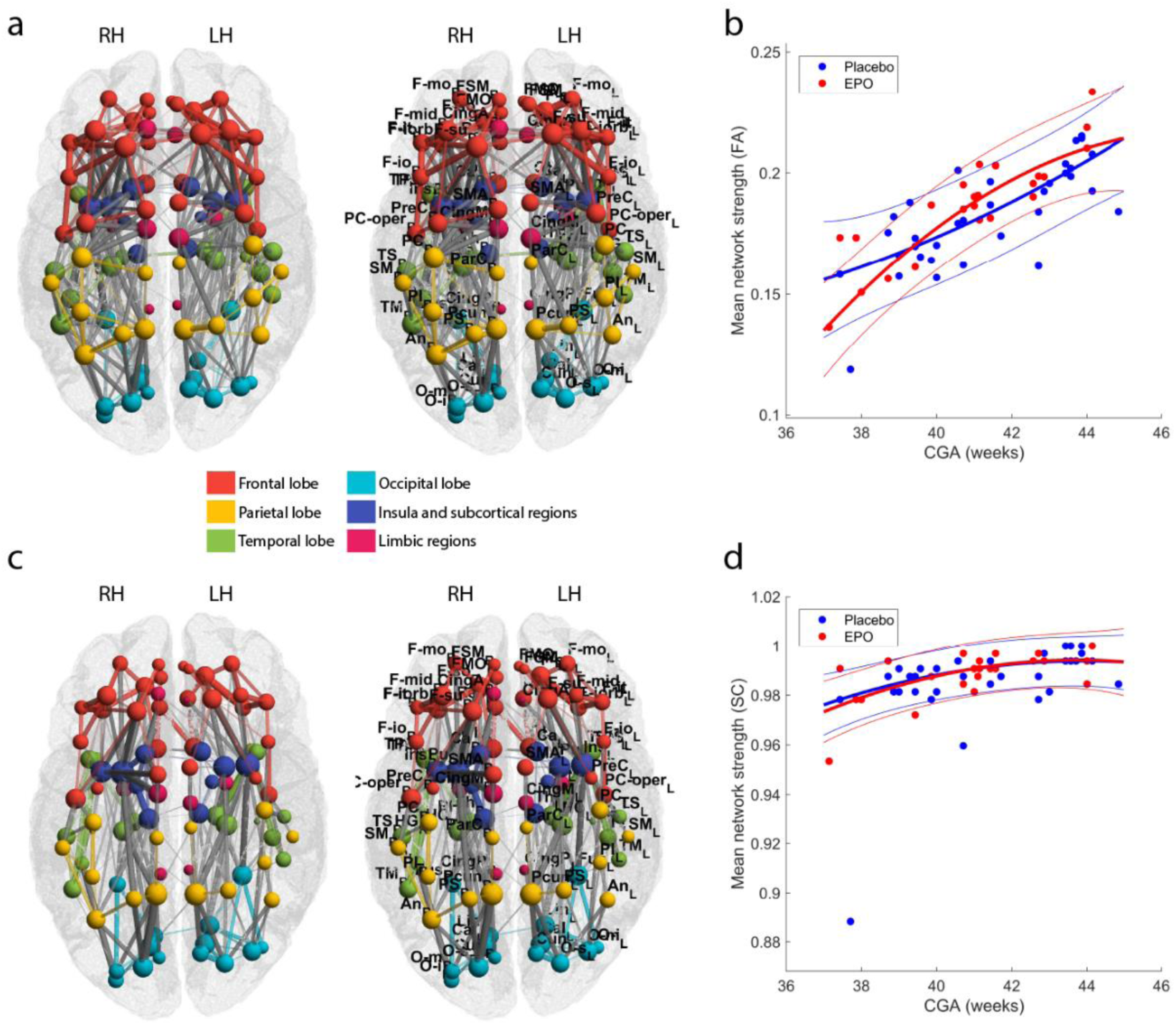
Age related changes of structural connectivity in the EPO-treated and placebo infant groups. *(a) Structural connectivity network where corrected gestational age at MRI (CGA) significantly positively correlated with the FA*_*mean*_ *of graph edges, (b) structural connectivity network where CGA was significantly correlated with the SC. For the abbreviations of ROIs, see Table S1.*

**Supplementary Figure S2.**
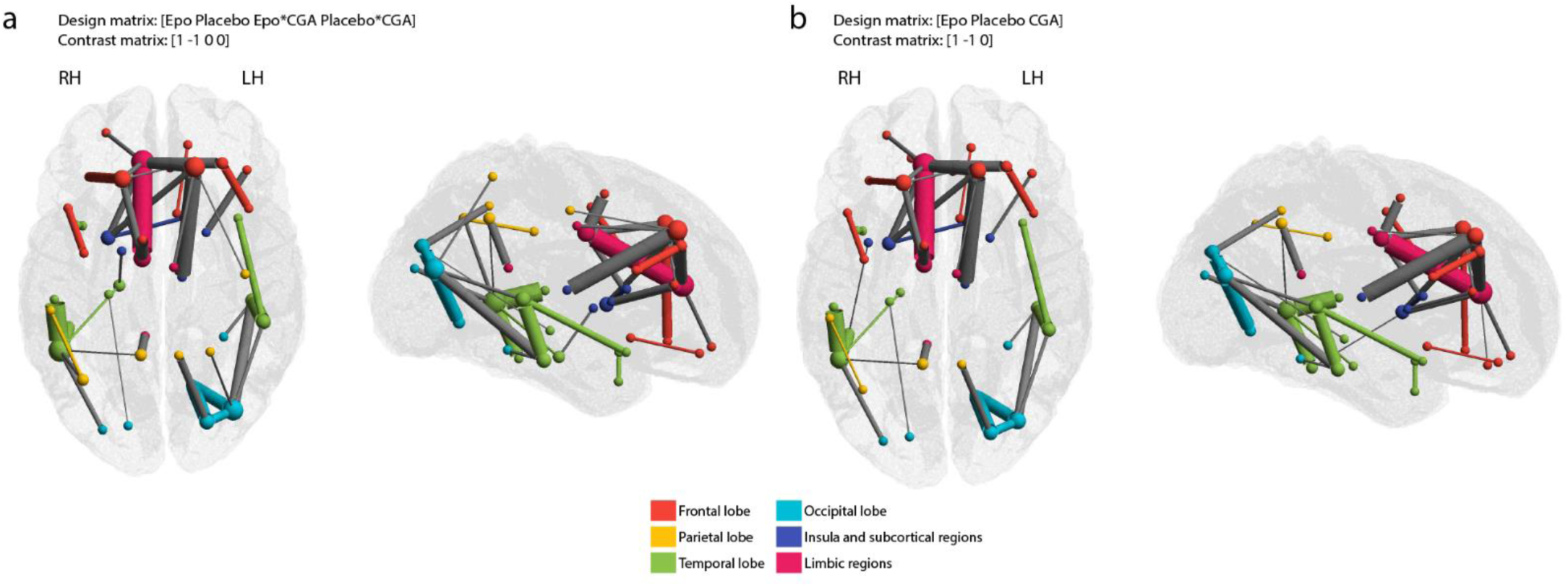
Analysis of the effect of EPO in the presence of group-CGA interaction. *The effect of EPO treatment on the FA*_*mean*_ *network is demon-strated (a) with Group-CGA interaction term and (b) without covariate interaction term in the model. For the SC network, the same analyses have been performed (c,d).*

**Supplementary Table 1.** Region of interest abbreviations and locations.

| Name of region | Short | Lobe / region | COG, x | COG, y | COG, z |
| --- | --- | --- | --- | --- | --- |
| Left precentral gyrus | PreC_L | Frontal | 24.5 | -7.0 | 37.3 |
| Right precentral gyrus | PreC_R | Frontal | -21.3 | -9.9 | 38.4 |
| Left superior frontal gyrus, dorsolateral | F-su_L | Frontal | 11.9 | 15.9 | 31.4 |
| Right superior frontal gyrus, dorsolateral | F-su_R | Frontal | -9.9 | 12.3 | 33.4 |
| Left superior frontal gyrus, orbital part | F-so_L | Frontal | 9.9 | 22.6 | -5.4 |
| Right superior frontal gyrus, orbital part | F-so_R | Frontal | -7.5 | 19.5 | -6.2 |
| Left middle frontal gyrus, lateral part | F-mid_L | Frontal | 20.1 | 17.1 | 26.7 |
| Right middle frontal gyrus, lateral part | F-mid_R | Frontal | -18.5 | 16.9 | 25.3 |
| Left middle frontal gyrus, orbital part | F-mo_L | Frontal | 17.6 | 26.1 | -2.1 |
| Right middle frontal gyrus, orbital part | F-mo_R | Frontal | -14.6 | 26.2 | -3.4 |
| Left opercular part of inferior frontal gyrus | F-io_L | Frontal | 28.1 | 3.5 | 19.4 |
| Right opercular part of inferior frontal gyrus | F-io_R | Frontal | -26.1 | 3.6 | 19.9 |
| Left area triangularis | F-it_L | Frontal | 27.6 | 15.1 | 15.1 |
| Right area triangularis | F-it_R | Frontal | -25.8 | 13.5 | 13.4 |
| Left orbital part of inferior frontal gyrus | F-iorb_L | Frontal | 20.8 | 13.4 | -1.6 |
| Right orbital part of inferior frontal gyrus | F-iorb_R | Frontal | -19.8 | 13.0 | -1.9 |
| Left rolandic operculum | PC-oper_L | Parietal | 28.2 | -11.8 | 19.5 |
| Right rolandic operculum | PC-oper_R | Parietal | -27.5 | -12.9 | 18.1 |
| Left supplementary motor area | SMA_L | Frontal | 5.5 | -3.9 | 44.2 |
| Right supplementary motor area | SMA_R | Frontal | -3.8 | -5.9 | 42.4 |
| Left olfactory cortex | OL_L | Frontal | 6.6 | 2.0 | -0.8 |
| Right olfactory cortex | OL_R | Frontal | -4.0 | 1.6 | -1.3 |
| Left superior frontal gyrus, medial part | FSM_L | Frontal | 5.1 | 24.6 | 23.8 |
| Right superior frontal gyrus, medial part | FSM_R | Frontal | -3.5 | 25.8 | 22.8 |
| Left superior frontal gyrus, medial orbital part | FMO_L | Frontal | 3.6 | 25.4 | -2.6 |
| Right superior frontal gyrus, medial orbital part | FMO_R | Frontal | -2.9 | 22.2 | -1.2 |
| Left gyrus rectus | Rec_L | Frontal | 4.9 | 16.1 | -7.6 |
| Right gyrus rectus | Rec_R | Frontal | -3.2 | 13.6 | -7.3 |
| Left insula | Ins_L | Subcortical/insula | 22.2 | -3.1 | 10.0 |
| Right insula | Ins_R | Subcortical/insula | -20.0 | -5.0 | 9.1 |
| Left anterior cingulate gyrus | CingA_L | Limbic | 4.6 | 15.7 | 14.2 |
| Right anterior cingulate gyrus | CingA_R | Limbic | -3.7 | 17.4 | 15.0 |
| Left middle cingulate | CingM_L | Limbic | 5.7 | -14.1 | 32.8 |
| Right middle cingulate | CingM_R | Limbic | -3.8 | -11.4 | 30.6 |
| Left posterior cingulate gyrus | CingP_L | Limbic | 5.2 | -33.4 | 22.2 |
| Right posterior cingulate gyrus | CingP_R | Limbic | -3.0 | -34.7 | 20.1 |
| Left hippocampus | HC_L | Temporal | 16.1 | -20.4 | 1.5 |
| Right hippocampus | HC_R | Temporal | -13.8 | -22.2 | 1.4 |
| Left parahippocampal gyrus | PH_L | Temporal | 13.3 | -17.8 | -6.6 |
| Right parahippocampal gyrus | PH_R | Temporal | -10.7 | -19.6 | -6.5 |
| Left amygdala | Am_L | Limbic | 15.8 | -8.3 | -4.1 |
| Right amygdala | Am_R | Limbic | -13.4 | -9.9 | -5.0 |
| Left calcarine sulcus | Cal_L | Occipital | 9.9 | -54.9 | 12.7 |
| Right calcarine sulcus | Cal_R | Occipital | -5.8 | -55.5 | 13.1 |
| Left cuneus | Cun_L | Occipital | 8.6 | -57.6 | 25.5 |
| Right cuneus | Cun_R | Occipital | -3.4 | -58.8 | 26.6 |
| Left lingual gyrus | Lin_L | Occipital | 11.1 | -49.5 | 3.8 |
| Right lingual gyrus | Lin_R | Occipital | -4.8 | -52.5 | 2.7 |
| Left superior occipital | O-s_L | Occipital | 15.5 | -60.4 | 26.7 |
| Right superior occipital | O-s_R | Occipital | -8.0 | -61.5 | 26.7 |
| Left middle occipital | O-m_L | Occipital | 23.4 | -56.9 | 19.8 |
| Right middle occipital | O-m_R | Occipital | -15.7 | -62.8 | 19.9 |
| Left inferior occipital | O-i_L | Occipital | 26.3 | -55.8 | 2.8 |
| Right inferior occipital | O-i_R | Occipital | -15.9 | -65.3 | 0.7 |
| Left fusiform gyrus | Fus_L | Occipital | 20.6 | -34.9 | -4.4 |
| Right fusiform gyrus | Fus_R | Occipital | -15.5 | -37.6 | -4.9 |
| Left postcentral gyrus | PC_L | Parietal | 27.0 | -16.2 | 37.4 |
| Right postcentral gyrus | PC_R | Parietal | -22.4 | -19.7 | 38.7 |
| Left superior parietal lobule | PS_L | Parietal | 16.6 | -39.4 | 47.5 |
| Right superior parietal lobule | PS_R | Parietal | -10.7 | -41.9 | 48.7 |
| Left inferior parietal lobule | PI_L | Parietal | 27.0 | -31.4 | 40.9 |
| Right inferior parietal lobule | PI_R | Parietal | -22.7 | -35.4 | 40.5 |
| Left supramarginal gyrus | SM_L | Parietal | 35.1 | -23.8 | 31.2 |
| Right supramarginal gyrus | SM_R | Parietal | -31.3 | -27.1 | 31.1 |
| Left angular gyrus | An_L | Parietal | 29.2 | -42.1 | 34.4 |
| Right angular gyrus | An_R | Parietal | -21.6 | -47.8 | 34.8 |
| Left precuneus | Pcun_L | Parietal | 7.1 | -40.5 | 38.7 |
| Right precuneus | Pcun_R | Parietal | -4.3 | -40.4 | 35.0 |
| Left paracentral lobule | ParC_L | Parietal | 5.7 | -21.3 | 48.8 |
| Right paracentral lobule | ParC_R | Parietal | -4.1 | -24.2 | 46.6 |
| Left caudate nucleus | Ca_L | Subcortical/insula | 8.8 | 0.7 | 13.8 |
| Right caudate nucleus | Ca_R | Subcortical/insula | -7.6 | 0.0 | 13.9 |
| Left putamen | Pu_L | Subcortical/insula | 15.4 | -3.7 | 10.1 |
| Right putamen | Pu_R | Subcortical/insula | -14.0 | -5.2 | 9.3 |
| Left globus pallidus | Pa_L | Subcortical/insula | 11.9 | -8.0 | 9.3 |
| Right globus pallidus | Pa_R | Subcortical/insula | -9.9 | -9.2 | 8.1 |
| Left thalamus | Th_L | Subcortical/insula | 8.1 | -17.1 | 13.5 |
| Right thalamus | Th_R | Subcortical/insula | -6.4 | -18.7 | 13.5 |
| Left transverse temporal gyri | HG_L | Temporal | 24.9 | -19.4 | 16.1 |
| Right transverse temporal gyri | HG_R | Temporal | -22.6 | -22.1 | 15.3 |
| Left superior temporal gyrus | TS_L | Temporal | 32.7 | -18.0 | 14.6 |
| Right superior temporal gyrus | TS_R | Temporal | -30.1 | -23.7 | 13.4 |
| Left superior temporal pole | TPS_L | Temporal | 24.9 | 0.6 | -5.1 |
| Right superior temporal pole | TPS_R | Temporal | -23.5 | -1.5 | -5.1 |
| Left middle temporal gyrus | TM_L | Temporal | 32.5 | -30.4 | 10.8 |
| Right middle temporal gyrus | TM_R | Temporal | -28.4 | -38.9 | 9.8 |
| Left middle temporal pole | TPM_L | Temporal | 23.3 | 0.9 | -14.0 |
| Right middle temporal pole | TPM_R | Temporal | -21.7 | -1.6 | -13.9 |
| Left inferior temporal gyrus | TI_L | Temporal | 29.3 | -24.1 | -7.5 |
| Right inferior temporal gyrus | TI_R | Temporal | -25.7 | -31.6 | -7.0 |

## References

Aja-Fernandez, S., Tristan-Vega, A., Alberola-Lopez, C., 2009. Noise estimation in single- and multiple-coil magnetic resonance data based on statistical models. Magn. Reson. Imaging. 27, 1397–1409.

Andersson, J.L.R., Sotiropoulos, S.N., 2016. An integrated approach to correction for off-resonance effects and subject movement in diffusion MR imaging. Neuroimage. 125, 1063–1078.

Back, S.A., Riddle, A., McClure, M.M., 2007. Maturation-dependent vulnerability of peri-natal white matter in premature birth. Stroke. 38, 724–730.

Ball, G., Aljabar, P., Zebari, S., Tusor, N., Arichi, T., Merchant, N., Robinson, E.C., Ogundipe, E., Rueckert, D., Edwards, A.D., Counsell, S.J., 2014. Rich-club organization of the newborn human brain. Proc. Natl. Acad. Sci. U. S. A. 111, 7456–7461.

Berman, J.I., Mukherjee, P., Partridge, S.C., Miller, S.P., Ferriero, D.M., Barkovich, A.J., Vigneron, D.B., Henry, R.G., 2005. Quantitative diffusion tensor MRI fiber tractography of sensorimotor white matter development in premature infants. Neuroimage. 27, 862–871.

Bonifacio, S.L., Glass, H.C., Chau, V., Berman, J.I., Xu, D., Brant, R., Barkovich, A.J., Poskitt, K.J., Miller, S.P., Ferriero, D.M., 2010. Extreme premature birth is not associated with impaired development of brain microstructure. J. Pediatr. 157, 726-32.e1.

Bonilha, L., Jensen, J.H., Baker, N., Breedlove, J., Nesland, T., Lin, J.J., Drane, D.L., Saindane, A.M., Binder, J.R., Kuzniecky, R.I., 2015. The brain connectome as a personalized biomarker of seizure outcomes after temporal lobectomy. Neurology. 84, 1846–1853.

Borgatti, S.P., Everett, M.G., 2000. Models of core/periphery structures. Social Networks. 21, 375–395.

Brown, C.J., Miller, S.P., Booth, B.G., Andrews, S., Chau, V., Poskitt, K.J., Hamarneh, G., 2014. Structural network analysis of brain development in young preterm neonates. Neuroimage. 101, 667–680.

Brydges, C.R., Landes, J.K., Reid, C.L., Campbell, C., French, N., Anderson, M., 2018. Cognitive outcomes in children and adolescents born very preterm: A meta-analysis. Dev. Med. Child Neurol. 60, 452–468.

Cao, M., Huang, H., He, Y., 2017. Developmental connectomics from infancy through early childhood. Trends Neurosci. 40, 494–506.

Collette, F., Hogge, M., Salmon, E., Van der Linden, M., 2006. Exploration of the neural substrates of executive functioning by functional neuroimaging. Neuroscience. 139, 209–221.

Crossley, N.A., Marques, T.R., Taylor, H., Chaddock, C., Dell’Acqua, F., Reinders, A.A., Mondelli, V., DiForti, M., Simmons, A., David, A.S., Kapur, S., Pariante, C.M., Murray, R.M., Dazzan, P., 2017. Connectomic correlates of response to treatment in first-episode psychosis. Brain. 140, 487–496.

de Reus, M.A., van den Heuvel, M.P., 2013. Estimating false positives and negatives in brain networks. Neuroimage. 70, 402–409.

Doria, V., Beckmann, C.F., Arichi, T., Merchant, N., Groppo, M., Turkheimer, F.E., Counsell, S.J., Murgasova, M., Aljabar, P., Nunes, R.G., Larkman, D.J., Rees, G., Edwards, A.D., 2010. Emergence of resting state networks in the preterm human brain. Proc. Natl. Acad. Sci. U. S. A. 107, 20015–20020.

Dosenbach, N.U., Fair, D.A., Miezin, F.M., Cohen, A.L., Wenger, K.K., Dosenbach, R.A., Fox, M.D., Snyder, A.Z., Vincent, J.L., Raichle, M.E., Schlaggar, B.L., Petersen, S.E., 2007. Distinct brain networks for adaptive and stable task control in humans. Proc. Natl. Acad. Sci. U. S. A. 104, 11073–11078.

Fauchere, J.C., Dame, C., Vonthein, R., Koller, B., Arri, S., Wolf, M., Bucher, H.U., 2008. An approach to using recombinant erythropoietin for neuroprotection in very preterm infants. Pediatrics. 122, 375–382.

Fischer, H.S., Reibel, N.J., Buhrer, C., Dame, C., 2017. Prophylactic early erythropoietin for neuroprotection in preterm infants: A meta-analysis. Pediatrics. 139, 10.1542/peds.2016-4317. Epub 2017 Apr 7.

Fischi-Gomez, E., Vasung, L., Meskaldji, D.E., Lazeyras, F., Borradori-Tolsa, C., Hagmann, P., Barisnikov, K., Thiran, J.P., Huppi, P.S., 2015. Structural brain connectivity in school-age preterm infants provides evidence for impaired networks relevant for higher order cognitive skills and social cognition. Cereb. Cortex. 25, 2793–2805.

Fischi-Gomez, E., Munoz-Moreno, E., Vasung, L., Griffa, A., Borradori-Tolsa, C., Monnier, M., Lazeyras, F., Thiran, J.P., Huppi, P.S., 2016. Brain network characterization of high-risk preterm-born school-age children. Neuroimage Clin. 11, 195–209.

Freeman, L.C., 1979. Centrality in social networks conceptual clarification. Social Networks. 1, 215–239.

Gonzalez, F.F., Abel, R., Almli, C.R., Mu, D., Wendland, M., Ferriero, D.M., 2009. Erythropoietin sustains cognitive function and brain volume after neonatal stroke. Dev. Neurosci. 31, 403–411.

Hagmann, P., Grant, P.E., Fair, D.A., 2012. MR connectomics: A conceptual framework for studying the developing brain. Front. Syst. Neurosci. 6, 43.

Hagmann, P., Cammoun, L., Gigandet, X., Meuli, R., Honey, C.J., Wedeen, V.J., Sporns, O., 2008. Mapping the structural core of human cerebral cortex. PLoS Biol. 6, e159.

Haynes, R.L., Folkerth, R.D., Keefe, R.J., Sung, I., Swzeda, L.I., Rosenberg, P.A., Volpe, J.J., Kinney, H.C., 2003. Nitrosative and oxidative injury to premyelinating oligodendrocytes in periventricular leukomalacia. J. Neuropathol. Exp. Neurol. 62, 441–450.

Huppi, P.S., Maier, S.E., Peled, S., Zientara, G.P., Barnes, P.D., Jolesz, F.A., Volpe, J.J., 1998. Microstructural development of human newborn cerebral white matter assessed in vivo by diffusion tensor magnetic resonance imaging. Pediatr. Res. 44, 584–590.

Jakab, A., Kasprian, G., Schwartz, E., Gruber, G.M., Mitter, C., Prayer, D., Schopf, V., Langs, G., 2015. Disrupted developmental organization of the structural connectome in fetuses with corpus callosum agenesis. Neuroimage. 111, 277–288.

Jurado, M.B., Rosselli, M., 2007. The elusive nature of executive functions: A review of our current understanding. Neuropsychol. Rev. 17, 213–233.

Juul, S., 2012. Neuroprotective role of erythropoietin in neonates. J. Matern. Fetal. Neonatal Med. 25 Suppl 4, 105–107.

Juul, S., 2010. Erythropoietin as a neonatal neuroprotective agent. NeoReviews. 11, e78.

Karolis, V.R., Froudist-Walsh, S., Brittain, P.J., Kroll, J., Ball, G., Edwards, A.D., Dell’Acqua, F., Williams, S.C., Murray, R.M., Nosarti, C., 2016. Reinforcement of the brain’s rich-club architecture following early neurodevelopmental disruption caused by very preterm birth. Cereb. Cortex. 26, 1322–1335.

Kinney, H.C., Brody, B.A., Kloman, A.S., Gilles, F.H., 1988. Sequence of central nervous system myelination in human infancy. II. patterns of myelination in autopsied infants. J. Neuropathol. Exp. Neurol. 47, 217–234.

Kuhn, T., Gullett, J.M., Nguyen, P., Boutzoukas, A.E., Ford, A., Colon-Perez, L.M., Triplett, W., Carney, P.R., Mareci, T.H., Price, C.C., Bauer, R.M., 2015. Test-retest reliability of high angular resolution diffusion imaging acquisition within medial temporal lobe connections assessed via tract based spatial statistics, probabilistic tractography and a novel graph theory metric. Brain Imaging Behav.

Latal, B., 2009. Prediction of neurodevelopmental outcome after preterm birth. Pediatr. Neurol. 40, 413–419.

Lebel, C., Benner, T., Beaulieu, C., 2012. Six is enough? comparison of diffusion parameters measured using six or more diffusion-encoding gradient directions with deterministic tractography. Magn. Reson. Med. 68, 474–483.

Leuchter, R.H., Gui, L., Poncet, A., Hagmann, C., Lodygensky, G.A., Martin, E., Koller, B., Darque, A., Bucher, H.U., Huppi, P.S., 2014. Association between early administration of high-dose erythropoietin in preterm infants and brain MRI abnormality at term-equivalent age. Jama. 312, 817–824.

Natalucci, G., Latal, B., Koller, B., Ruegger, C., Sick, B., Held, L., Bucher, H.U., Fauchere, J.C., Swiss EPO Neuroprotection Trial Group, 2016. Effect of early prophylactic high-dose recombinant human erythropoietin in very preterm infants on neurodevelopmental outcome at 2 years: A randomized clinical trial. Jama. 315, 2079–2085.

Newman, M.E., 2006. Finding community structure in networks using the eigenvectors of matrices. Phys. Rev. E. Stat. Nonlin Soft Matter Phys. 74, 036104.

O’Gorman, R.L., Bucher, H.U., Held, U., Koller, B.M., Huppi, P.S., Hagmann, C.F., Swiss EPO Neuroprotection Trial Group, 2015. Tract-based spatial statistics to assess the neuroprotective effect of early erythropoietin on white matter development in pre-term infants. Brain. 138, 388–397.

Onnela, J., Saram\aki, J., Kert\’esz, J., Kaski, K., 2005. Intensity and coherence of motifs in weighted complex networks. Phys Rev E. 71, 065103.

Pandit, A.S., Robinson, E., Aljabar, P., Ball, G., Gousias, I.S., Wang, Z., Hajnal, J.V., Rueckert, D., Counsell, S.J., Montana, G., Edwards, A.D., 2014. Whole-brain mapping of structural connectivity in infants reveals altered connection strength associated with growth and preterm birth. Cereb. Cortex. 24, 2324–2333.

Parker, G.J., Haroon, H.A., Wheeler-Kingshott, C.A., 2003. A framework for a streamline-based probabilistic index of connectivity (PICo) using a structural interpretation of MRI diffusion measurements. J. Magn. Reson. Imaging. 18, 242–254.

Reess, T.J., Rus, O.G., Schmidt, R., de Reus, M.A., Zaudig, M., Wagner, G., Zimmer, C., van den Heuvel, M.P., Koch, K., 2016. Connectomics-based structural network alterations in obsessive-compulsive disorder. Transl. Psychiatry. 6, e882.

Riddle, A., Luo, N.L., Manese, M., Beardsley, D.J., Green, L., Rorvik, D.A., Kelly, K.A., Barlow, C.H., Kelly, J.J., Hohimer, A.R., Back, S.A., 2006. Spatial heterogeneity in oligodendrocyte lineage maturation and not cerebral blood flow predicts fetal ovine periventricular white matter injury. J. Neurosci. 26, 3045–3055.

Robertson, N.J., Tan, S., Groenendaal, F., van Bel, F., Juul, S.E., Bennet, L., Derrick, M., Back, S.A., Valdez, R.C., Northington, F., Gunn, A.J., Mallard, C., 2012. Which neuro-protective agents are ready for bench to bedside translation in the newborn infant? J. Pediatr. 160, 544-552.e4.

Rose, S.E., Hatzigeorgiou, X., Strudwick, M.W., Durbridge, G., Davies, P.S., Colditz, P.B., 2008. Altered white matter diffusion anisotropy in normal and preterm infants at term-equivalent age. Magn. Reson. Med. 60, 761–767.

Rubinov, M., Sporns, O., 2010. Complex network measures of brain connectivity: Uses and interpretations. Neuroimage. 52, 1059–1069.

Ruegger, C.M., Kraus, A., Koller, B., Natalucci, G., Latal, B., Waldesbuhl, E., Fauchere, J.C., Held, L., Bucher, H.U., 2014. Randomized controlled trials in very preterm infants: Does inclusion in the study result in any long-term benefit? Neonatology. 106, 114– 119.

Saigal, S., Doyle, L.W., 2008. An overview of mortality and sequelae of preterm birth from infancy to adulthood. Lancet. 371, 261–269.

Shi, F., Yap, P.T., Wu, G., Jia, H., Gilmore, J.H., Lin, W., Shen, D., 2011. Infant brain atlases from neonates to 1- and 2-year-olds. PLoS One. 6, e18746.

Shingo, T., Sorokan, S.T., Shimazaki, T., Weiss, S., 2001. Erythropoietin regulates the in vitro and in vivo production of neuronal progenitors by mammalian forebrain neural stem cells. J. Neurosci. 21, 9733–9743.

Sporns, O., 2011. The human connectome: A complex network. Ann. N. Y. Acad. Sci. 1224, 109–125.

Sporns, O., 2013. Network attributes for segregation and integration in the human brain. Curr. Opin. Neurobiol. 23, 162–171.

Sugawa, M., Sakurai, Y., Ishikawa-Ieda, Y., Suzuki, H., Asou, H., 2002. Effects of erythropoietin on glial cell development; oligodendrocyte maturation and astrocyte proliferation. Neurosci. Res. 44, 391–403.

Tsai, P.T., Ohab, J.J., Kertesz, N., Groszer, M., Matter, C., Gao, J., Liu, X., Wu, H., Carmichael, S.T., 2006. A critical role of erythropoietin receptor in neurogenesis and post-stroke recovery. J. Neurosci. 26, 1269–1274.

Tymofiyeva, O., Hess, C.P., Ziv, E., Tian, N., Bonifacio, S.L., McQuillen, P.S., Ferriero, D.M., Barkovich, A.J., Xu, D., 2012. Towards the “baby connectome”: Mapping the structural connectivity of the newborn brain. PLoS One. 7, e31029.

Tzourio-Mazoyer, N., Landeau, B., Papathanassiou, D., Crivello, F., Etard, O., Delcroix, N., Mazoyer, B., Joliot, M., 2002. Automated anatomical labeling of activations in SPM using a macroscopic anatomical parcellation of the MNI MRI single-subject brain. Neuroimage. 15, 273–289.

van den Heuvel, M.P., Sporns, O., 2011. Rich-club organization of the human connectome. J. Neurosci. 31, 15775–15786.

van den Heuvel, M.P., Sporns, O., 2013. An anatomical substrate for integration among functional networks in human cortex. J. Neurosci. 33, 14489–14500.

van den Heuvel, M.P., Kersbergen, K.J., de Reus, M.A., Keunen, K., Kahn, R.S., Groenendaal, F., de Vries, L.S., Benders, M.J., 2015. The neonatal connectome during preterm brain development. Cereb. Cortex. 25, 3000–3013.

Vollmer, B., Lundequist, A., Martensson, G., Nagy, Z., Lagercrantz, H., Smedler, A.C., Forssberg, H., 2017. Correlation between white matter microstructure and executive functions suggests early developmental influence on long fibre tracts in preterm born adolescents. PLoS One. 12, e0178893.

Volpe, J.J., 2009. Brain injury in premature infants: A complex amalgam of destructive and developmental disturbances. Lancet Neurol. 8, 110–124.

Wakana, S., Caprihan, A., Panzenboeck, M.M., Fallon, J.H., Perry, M., Gollub, R.L., Hua, K., Zhang, J., Jiang, H., Dubey, P., Blitz, A., van Zijl, P., Mori, S., 2007. Reproducibility of quantitative tractography methods applied to cerebral white matter. Neuroimage. 36, 630–644.

Watts, D.J., Strogatz, S.H., 1998. Collective dynamics of ‘small-world’ networks. Nature. 393, 440–442.

Woodward, L.J., Anderson, P.J., Austin, N.C., Howard, K., Inder, T.E., 2006. Neonatal MRI to predict neurodevelopmental outcomes in preterm infants. N. Engl. J. Med. 355, 685–694.

Woodward, L.J., Clark, C.A., Pritchard, V.E., Anderson, P.J., Inder, T.E., 2011. Neonatal white matter abnormalities predict global executive function impairment in children born very preterm. Dev. Neuropsychol. 36, 22–41.

Xia, M., Wang, J., He, Y., 2013. BrainNet viewer: A network visualization tool for human brain connectomics. PLoS One. 8, e68910.

Yung, A., Poon, G., Qiu, D.Q., Chu, J., Lam, B., Leung, C., Goh, W., Khong, P.L., 2007. White matter volume and anisotropy in preterm children: A pilot study of neurocognitive correlates. Pediatr. Res. 61, 732–736.

Zalesky, A., Fornito, A., Bullmore, E.T., 2010. Network-based statistic: Identifying differences in brain networks. Neuroimage. 53, 1197–1207.

Zeng, J., Luo, Q., Du, L., Liao, W., Li, Y., Liu, H., Liu, D., Fu, Y., Qiu, H., Li, X., Qiu, T., Meng, H., 2015. Reorganization of anatomical connectome following electroconvulsive therapy in major depressive disorder. Neural Plast. 2015, 271674.

